# Relapsing fever *Borrelia puertoricensis* in migratory Mexican free-tailed bats, Oklahoma, USA, 2022–2023

**DOI:** 10.64898/2026.01.13.699129

**Authors:** Daniel J. Becker, Kristin E. Dyer, Beckett L. Olbrys, Mackenzie G. Hightower, Meagan Allira, Bret Demory, Lauren R. Lock, Kiersten N. Taylor, Nakib N. Bhata, Selena M. Hernandez, Paul A. Lawson, Noha H. Youssef, Samuel L. Miller, Mostafa S. Elshahed, Taylor B. Verrett, Kerry L. Clark

## Abstract

We detected *Borrelia puertoricensis* in migratory Mexican free-tailed bats sampled in Oklahoma during 2022 and 2023, representing only the second detection of this relapsing fever species in wild vertebrates. Although prevalence was low (0.79%), our findings suggest migratory bats could contribute to dispersal of tick-borne pathogens in North America.

## Introduction

Bacteria in the genus *Borrelia* are primarily transmitted by ticks and cause two diseases of high human health concern: Lyme borreliosis (LB) and relapsing fever (RF). LB is the most common vector-borne disease in the Northern Hemisphere, with hard ticks (Ixodidae) serving as vectors of species in the *Borrelia burgdorferi* sensu lato complex (1). RF is globally distributed, with this clade of febrile illness–causing borreliae vectored by both hard and soft ticks (Argasidae) (2). Nymphal and adult ticks are the life stage most likely to bite humans and transmit borreliae, such that understanding which host species first infect tick larvae is critical to predicting human risk.

While rodents and migratory birds have long been understood to be important hosts in the epidemiology of LB and RF borreliae (2,3), bats have received comparatively less attention for their role in enzootic cycles and dispersal of these bacteria. However, bats can host RF borreliae that likely transmit to humans (4), and other borreliae hosted by bats in the tropics form novel clades adjacent to LB borreliae (5,6). Despite this growing attention, the genetic diversity and zoonotic potential of bat-borne borreliae remains poorly understood, in part because the vast majority of work has focused on bats in the Neotropics, Afrotropics, or Australasian realm (7).

Fewer bat-borne *Borrelia* surveys have been conducted in temperate zones (8), which could underestimate zoonotic potential and may ignore a key aspect of *Borrelia* transmission. Most migratory bats occur in temperate zones, and physiological costs of migration in birds can cause chronic *Borrelia* infection to reactivate (9). Migration could thus allow bats to not only disperse ticks but also active *Borrelia* infections. Here, we assessed *Borrelia* diversity in the Mexican free-tailed bat (*Tadarida brasiliensis*), which undertakes migrations between wintering grounds in Mexico and large, summer roosts throughout the southwestern United States (10).

### The Study

Within a larger study of bat migration, immunity, and infection (11), we collected blood from 386 Mexican free-tailed bats in western Oklahoma at monthly or biweekly intervals between March and October 2022 and 2023. We also opportunistically sampled pallid bats (*Antrozous pallidus, n* = 2), big brown bats (*Eptesicus fuscus, n* = 2), cave myotis (*Myotis velifer, n* = 11), and one hoary bat (*Lasiurus cinereus*). We captured bats with hand nets or mist nets and stored blood on Whatman FTA cards. Most of these 402 bats (75%) were inspected for ticks, with a subset collected and stored in 70% ethanol; we screened an additional four pallid bats, three western big-eared bats (*Corynorhinus townsendii*), and 21 Mexican free-tailed bats for ticks but did not collect blood (*n* = 427 screened). All bats were released at their capture site. Fieldwork was approved by the University of Oklahoma Institutional Animal Care and Use Committee (2022-0198), with permits from the Oklahoma Department of Wildlife Conservation (10567389).

We used QIAamp DNA Investigator Kits (Qiagen) to extract DNA from bat blood (11). Where possible, we microscopically identified collected ticks and extracted DNA using proteinase K digestion and *Quick*-DNA Miniprep Kits (Zymo). We confirmed tick taxonomy through PCR and Sanger sequencing of the *COI* gene (12). To determine *Borrelia* spp. infection, we used PCR and Sanger sequencing of the 16S rRNA and flagellin (*flaB*) genes (Table S1) (5). We aligned our sequences with references using MUSCLE and used MrBayes for phylogenetic analysis, with each gene tree run for 10,000,000 generations using a GTR + I + G model.

Across our two-year seasonal study, four bats were positive for borreliae (1%, 95% CI: 0.39–2.53%), including three Mexican free-tailed bats and one pallid bat (Table 1). Only 6.14% of Mexican free-tailed bats were parasitized by ticks, while we found ticks on 33% and 30% of pallid bats and cave myotis, respectively. No other bat species hosted borreliae or ticks. For Mexican free-tailed bats, parasitized hosts had one (72%), two (24%), or three (4%) ticks. We confirmed ticks as argasids (e.g., see GenBank PX651872), but none of three tested ticks were positive for borreliae. We then used generalized additive models to assess seasonal, annual, and sex variation in *Borrelia* infection and tick parasitism in Mexican free-tailed bats using cyclic smooth terms for week and the *mgcv* R package. Infection risk did not vary by week, year, or sex, while bat ticks were more likely in 2023 and after spring migration (Figure 1, Table S2). *Borrelia* and tick data have been deposited within the Pathogen Harmonized Observatory.

**Table 1.**
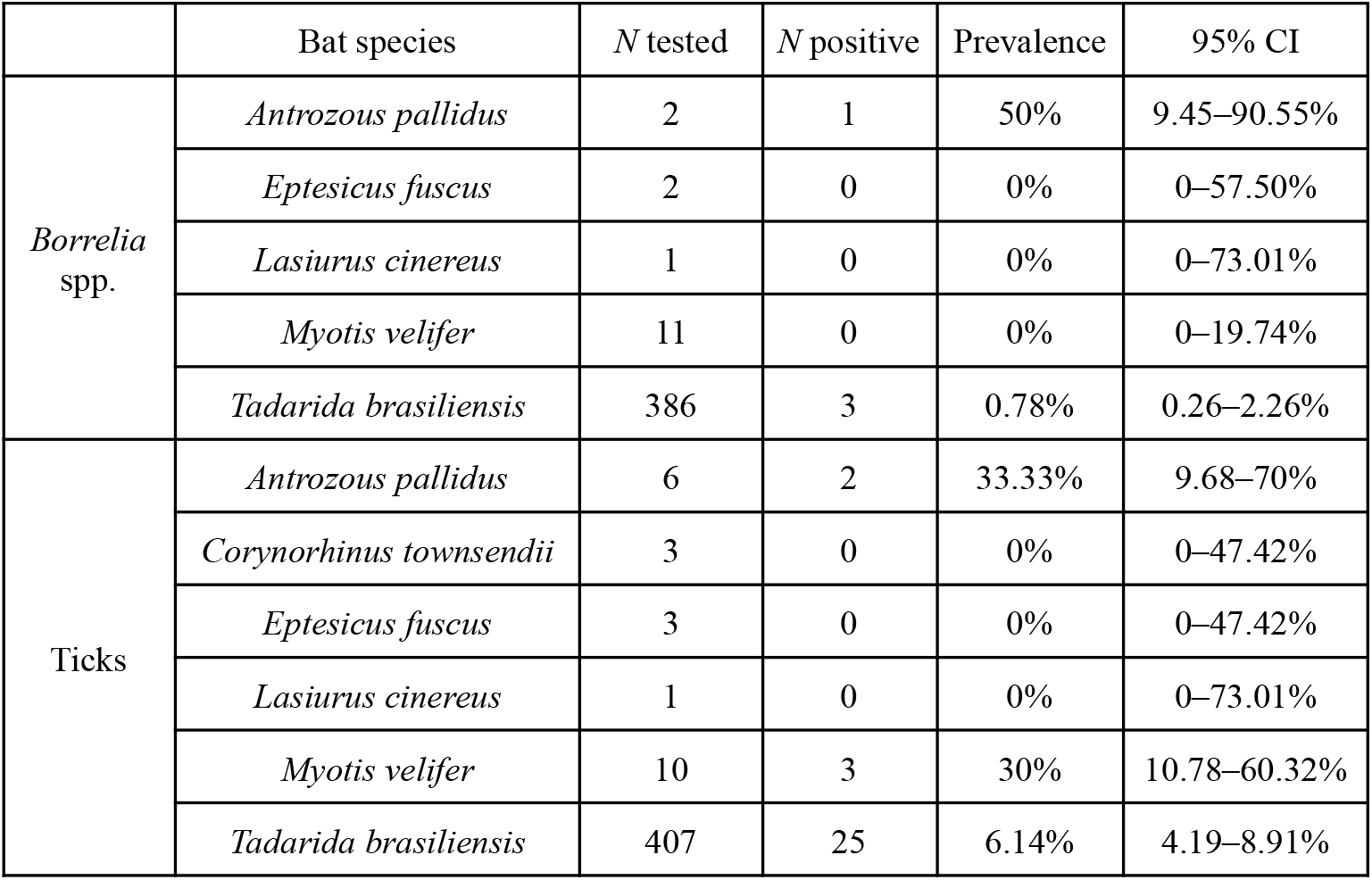
Bat species–level prevalence and 95% CIs (Wilson method) for *Borrelia* spp. infection and tick infestation status.

|  | Bat species | *N* tested | *N* positive | Prevalence | 95% CI |
| --- | --- | --- | --- | --- | --- |
| *Borrelia*<br>spp. | *Antrozous pallidus* | 2 | 1 | 50% | 9.45–90.55% |
|  | *Eptesicus fuscus* | 2 | 0 | 0% | 0–57.50% |
|  | *Lasiurus cinereus* | 1 | 0 | 0% | 0–73.01% |
|  | *Myotis velifer* | 11 | 0 | 0% | 0–19.74% |
|  | *Tadarida brasiliensis* | 386 | 3 | 0.78% | 0.26–2.26% |
| Ticks | *Antrozous pallidus* | 6 | 2 | 33.33% | 9.68–70% |
|  | *Corynorhinus townsendii* | 3 | 0 | 0% | 0–47.42% |
|  | *Eptesicus fuscus* | 3 | 0 | 0% | 0–47.42% |
|  | *Lasiurus cinereus* | 1 | 0 | 0% | 0–73.01% |
|  | *Myotis velifer* | 10 | 3 | 30% | 10.78–60.32% |
|  | *Tadarida brasiliensis* | 407 | 25 | 6.14% | 4.19–8.91% |

**Figure 1.**
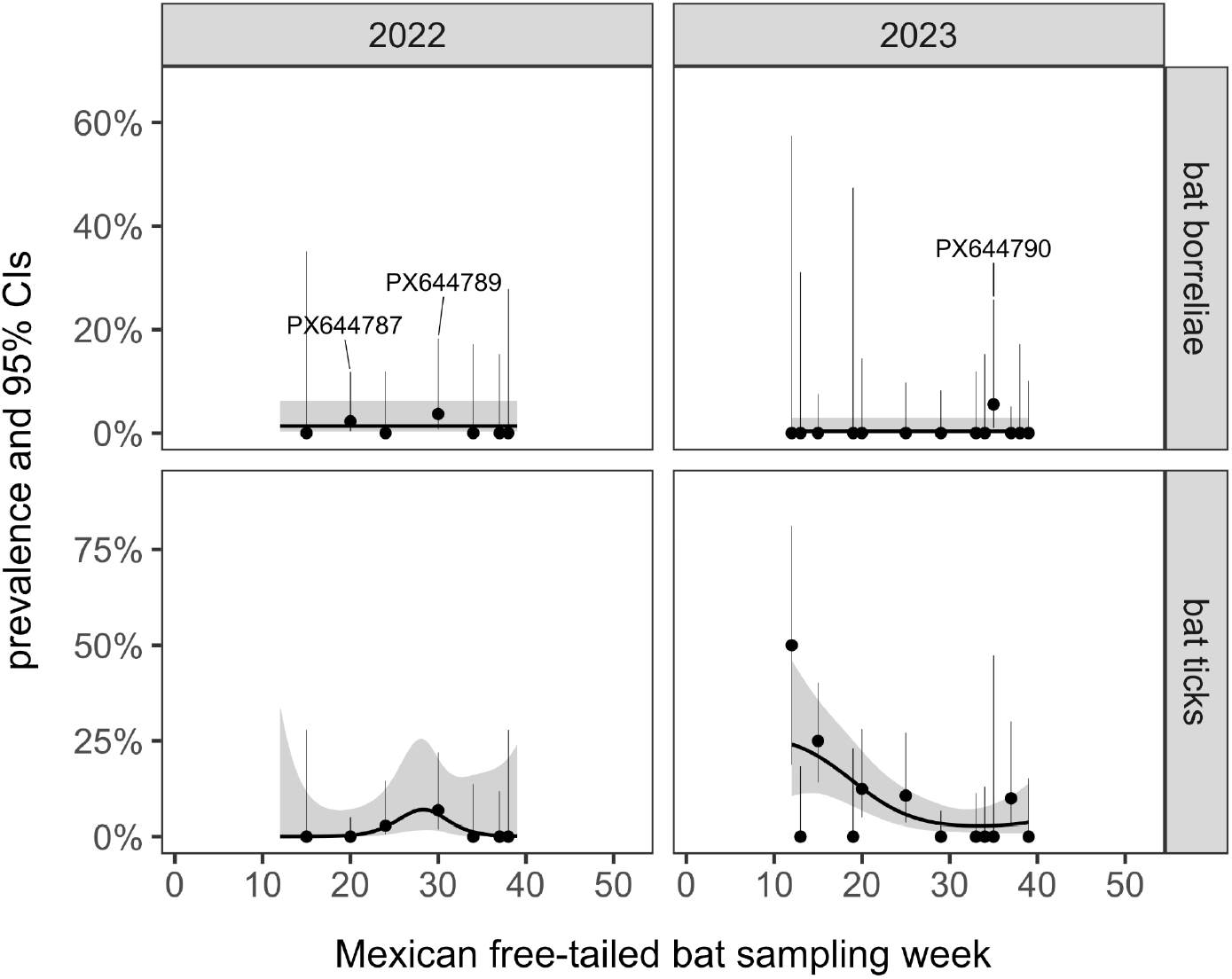
Seasonal and annual variation in the prevalence of *Borrelia* spp. infection and argasid tick parasitism in migratory Mexican free-tailed bats. Point estimates and 95% CIs (Wilson method) are provided alongside generalized additive model fitted values and 95% CIs. GenBank accessions for the 16S rRNA sequences of all three *Borrelia*-positive bats are matched to time.

We obtained 16S rRNA sequences from all positives, with *flaB* sequences from two of the Mexican free-tailed bats (Figure 2). Two 16S rRNA sequences from Mexican free-tailed bats showed 99.6–100% identity to *Borrelia puertoricensis* identified from soft ticks and opossums in the Neotropics (e.g., NR_181316) (13,14). The *flaB* sequences were also 97.5–98.4% similar to *Borrelia puertoricensis* (e.g., OQ944477), confirming these bat borreliae within the RF clade (Figure S1). Both positive bats were sampled just following or prior to migration. 16S rRNA sequences from the other Mexican free-tailed bat and the pallid bat formed two distinct lineages (92.8% identity). These were most related (91.1–95.2% similar) to novel borreliae found in *Pteropus* bats adjacent to the LB clade (5). Sequences are available on GenBank through accessions PX644787–90 (16S rRNA) and PX644764–65 (*flaB*).

**Figure 2.**
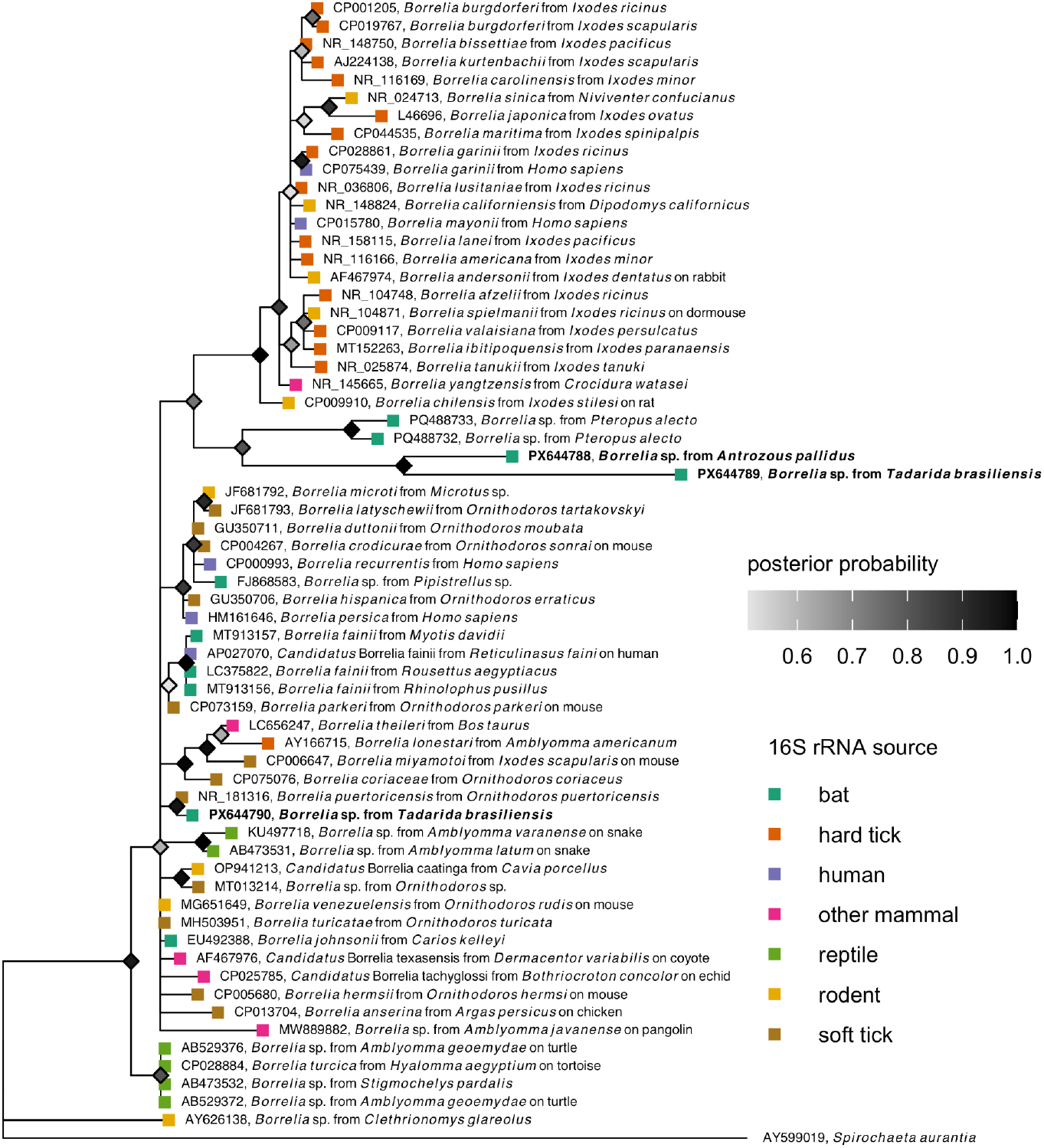
Consensus Bayesian phylogeny of the 16S rRNA *Borrelia* spp. sequences from this study (shown in bold) and reference sequences from bats, other mammals, reptiles, and ticks. Nodes are colored by posterior probability (nodes with less than 50% support are not shown).

Given the high similarity of 16S rRNA and *flaB* sequences to *Borrelia puertoricensis*, we attempted further characterization of the OK535 isolate using shotgun metagenomics (see Online Supplement). Given the low DNA volume, we first conducted whole genome amplification using multiple displacement amplification via the TruePrime WGA Kit (Expedeon) (15). Illumina sequencing yielded 17.48 Gbp in 58.07 million 300 bp paired-end reads, available via accessions PRJNA1401797 (BioProject), SAMN54571938 (BioSample), and SRR36790398 (SRA). We identified 1,592 paired-end reads (0.003% of total reads) affiliated with the genus *Borrelia* (File S1). tBLASTx query of assembled contigs (File S2) identified 622 partial or complete protein-coding genes (Table S3). The absolute majority of genes displayed either identical (*n* = 139) or extremely high (>80% amino acid identity, *n* = 467) to *Borrelia puertoricensis* reference genomes from soft ticks and a human patient (Figure 3, Table S4). These results corroborate *Borrelia puertoricensis* as the only *Borrelia* species in the Mexican free-tailed bat sample.

**Figure 3.**
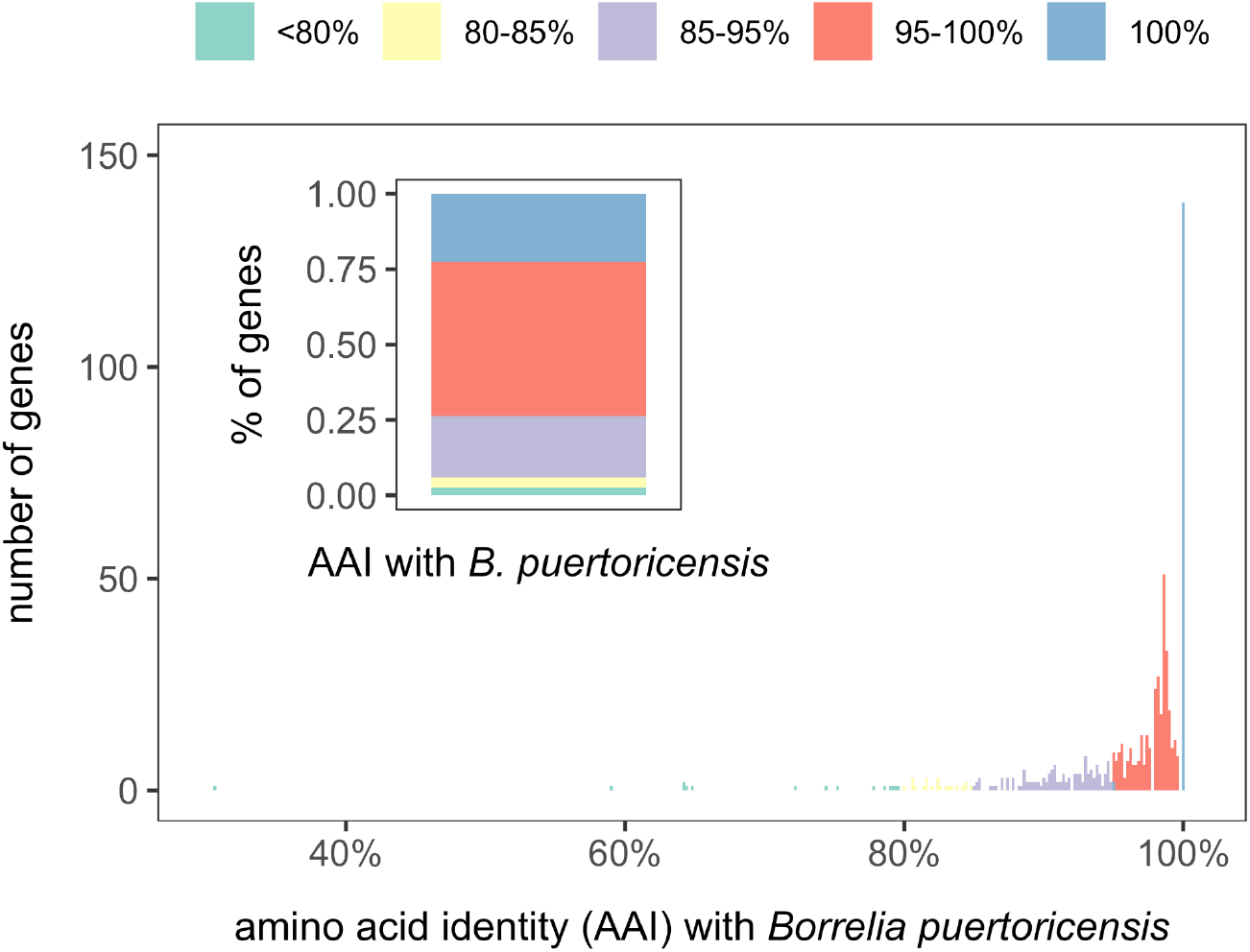
Amino acid identity (AAI) between the 622 *Borrelia* genes identified in the Mexican free-tailed bat sample (isolate OK535) and those from *Borrelia puertoricensis*, using reference genomes from soft ticks (GCA_023035875, NZ_CP149102) and a human (NZ_CP138334). See Table S3 and S4 for raw AAI data and all *Borrelia* genomes used in the bioinformatic analyses.

## Conclusions

We here demonstrate that migratory North American bats harbor distinct but LB-adjacent borreliae and RF infections. For the former, such results further support a novel clade of borreliae in bats spanning highly divergent host families (i.e., Pteropodidae, Molossidae, Vespertilionidae) (5). Future work is needed to characterize the zoonotic potential of these bat-hosted lineages. For RF infections, we identified *Borrelia puertoricensis* (13), representing the first detection of this lineage in bats and only the second detection in wild vertebrates (14). While these infections were rare, detection close to spring and fall migration could reflect the reactivation of chronic infections, as observed in birds (9). Similarly, such infections could indicate recent dispersal from wintering grounds, given prevalent tick infestations after spring migration and detection of *Borrelia puertoricensis* in soft ticks in Mexico (16). Our findings therefore warrant further attention to the role that migratory bats may play in the epidemiology and seasonal dispersal of RF borreliae as well as to the zoonotic risk of these infections.

## Acknowledgements

This work was supported by the National Institutes of Health (P20GM134973, P20GM152333), Research Corporation for Science Advancement (RCSA, Subawards 28365 and 29018, part of a USDA Non-Assistance Cooperative Agreement with RCSA Federal Award 58–3022–0-005), Edward Mallinckrodt, Jr. Foundation, National Science Foundation (DBI 2515340), and University of Oklahoma (Data Institute for Societal Challenges). We also thank Juliana Maria Nunes Batista and Molly Simonis for assistance with fieldwork.

## Biography

Dr. Becker is an Assistant Professor in the School of Biological Sciences at the University of Oklahoma. His work focuses on how zoonotic pathogens spread within and between populations and species and how environmental change alters infection dynamics, with an affinity for wild bat and songbird systems.

## References

1. Stone BL, Tourand Y, Brissette CA. Brave new worlds: The expanding universe of Lyme disease. Vector Borne Zoonotic Dis. 2017 Sep;17(9):619–29.

2. Jakab Á, Kahlig P, Kuenzli E, Neumayr A. Tick borne relapsing fever - a systematic review and analysis of the literature. PLoS Negl Trop Dis. 2022 Feb;16(2):e0010212.

3. Becker DJ, Han BA. The macroecology and evolution of avian competence for Borrelia burgdorferi. Glob Ecol Biogeogr. 2021 Mar;30(3):710–24.

4. Qiu Y, Nakao R, Hang’ombe BM, Sato K, Kajihara M, Kanchela S, et al. Human borreliosis caused by a New World relapsing fever Borrelia-like organism in the old world. Clin Infect Dis. 2019 Jun 18;69(1):107–12.

5. Verrett TB, Falvo CA, Benson E, Jones-Slobodian DN, Crowley DE, Dale AS, et al. Borrelia lineages adjacent to zoonotic clades in black flying foxes (Pteropus alecto), Australia, 2018-2020. Emerg Infect Dis. 2025 Jul;31(7):1415–20.

6. Muñoz-Leal S, Faccini-Martínez ÁA, Pérez-Torres J, Chala-Quintero SM, Herrera-Sepúlveda MT, Cuervo C, et al. Novel Borrelia genotypes in bats from the Macaregua Cave, Colombia. Zoonoses Public Health. 2021 Feb;68(1):12–8.

7. Szentivanyi T, McKee C, Jones G, Foster JT. Trends in Bacterial Pathogens of Bats: Global Distribution and Knowledge Gaps. Transbound Emerg Dis [Internet]. 2023 Mar 27 [cited 2023 Jul 10];2023. Available from: https://www.hindawi.com/journals/tbed/2023/9285855/

8. Banerjee A, Baid K, Byron T, Yip A, Ryan C, Thampy PR, et al. Seroprevalence in Bats and Detection of Borrelia burgdorferi in Bat Ectoparasites. Microorganisms. 2020 Mar 20;8(3):440.

9. Gylfe A, Bergström S, Lundström J, Olsen B. Reactivation of Borrelia infection in birds. Nature. 2000 Feb 17;403(6771):724–5.

10. Villa BR, Cockrum EL. Migration in the Guano Bat Tadarida Brasiliensis Mexicana (Saussure). J Mammal. 1962 Feb 28;43(1):43–64.

11. Becker DJ, Dyer KE, Lock LR, Olbrys BL, Pladas SA, Sukhadia AA, et al. Geographically widespread and novel hemotropic mycoplasmas and bartonellae in Mexican free-tailed bats and sympatric North American bat species. mSphere. 2024 Dec 11;e0011624.

12. Folmer O, Black MB, Hoeh W, Lutz R, Vrijenhoek R. DNA primers for amplification of mitochondrial cytochrome c oxidase subunit I from diverse metazoan invertebrates. Mol Mar Biol Biotechnol. 1994 Oct 1;3(5):294–9.

13. Bermúdez SE, Armstrong BA, Domínguez L, Krishnavajhala A, Kneubehl AR, Gunter SM, et al. Isolation and genetic characterization of a relapsing fever spirochete isolated from Ornithodoros puertoricensis collected in central Panama. PLoS Negl Trop Dis. 2021 Aug;15(8):e0009642.

14. López Y, Faccini-Martínez ÁA, Muñoz-Leal S, Contreras V, Calderón A, Rivero R, et al. Borrelia puertoricensis in opossums (Didelphis marsupialis) from Colombia. Parasit Vectors. 2023 Dec 4;16(1):448.

15. Long N, Qiao Y, Xu Z, Tu J, Lu Z. Recent advances and application in whole-genome multiple displacement amplification. Quant Biol. 2020 Dec;8(4):279–94.

16. Vázquez-Guerrero E, Kneubehl AR, Reyes-Solís GC, Machain-Williams C, Krishnavajhala A, Estrada-de Los Santos P, et al. Use of a mouse model for the isolation of Borrelia puertoricensis from soft ticks. PLoS One. 2025 Feb 18;20(2):e0318652.

