## Supplemental Material for "Relapsing fever *Borrelia puertoricensis* in migratory Mexican free-tailed bats, Oklahoma, USA, 2022–2023"

Table S1. PCR primers and reaction conditions used for *Borrelia* spp. Testing
Table S2. Results of generalized additive models for *Borrelia* spp. positivity and tick parasitism
Figure S1. Consensus Bayesian phylogeny of the two *flaB Borrelia* spp. sequences
Additional genomic methods
Table S3. BLAST results of all 622 partial or complete *Borrelia* genes identified
Table S4. List of reference *Borrelia* genomes used in bioinformatic analyses
File S1. FASTA files of reads affiliated with the genus *Borrelia*.
File S2. FASTA files of contigs assembled from *Borrelia* reads.

Table S1. PCR primers and reaction conditions used for *Borrelia* spp. testing*

| Gene | Primer | Orientation | Sequence, 5'-3' | Annealing (°C)** | Cycles | Product (bp) | Reference |
|---|---|---|---|---|---|---|---|
| 16S rRNA | 1A | F | CTAACGCTGGCAGTGCGTCTTAAGC | 70-61, 60 | 10 + 40 | ~724 | (1) |
| | 1B | R | AGCGTCAGTCTTGACCCAGAAGTTC | | | | |
| *flaB* | FlaLL | Outer F | ACATATTCAGATGCAGACAGAGG | 52 | 30 | ~665 | (2) |
| | FlaRL | Outer R | GCAATCATAGCCATTGCAGATTGT | | | | |
| . | 442f | Inner F | GCTGAAGAGCTTGGAATGCAACC | 55 | 30 | ~524 | (3) |
| | FlaRL | Inner R | GCAATCATAGCCATTGCAGATTGT | | | | (2) |

*PCRs used Promega GoTaq Green master mix. Primers were used at 0.5 mM concentration. All PCRs began with an initial denaturation step at 94°C, 2 minutes. Thereafter, cycles consisted of 94°C, 30 sec; annealing temp as indicated in the table for 30 sec; and extension at 72°C, 30 sec for 16S 1A/1B and *flaB* primers.

**The 16S rRNA screening primers (1A/1B) used a touchdown PCR approach. The annealing temperature was dropped one degree C in each of the first 10 cycles, followed by 40 additional cycles with annealing temperature as shown in the table.

References:


1. Richter D, Schlee DB, Matuschka FR. Relapsing fever-like spirochetes infecting European vector tick of Lyme disease agent. *Emerg Infect Dis*. 2003 Jun;9(6):697-701.
2. Barbour AG, Maupin GO, Teltow GJ, Carter CJ, Piesman J. Identification of an uncultivable *Borrelia* species in the hard tick *Amblyomma americanum*: possible agent of a Lyme disease-like illness. *J Infect Dis*. 1996 Feb;173(2):403-9.
3. Verrett TB, Falvo CA, Benson E, Jones-Slobodian DN, Crowley DE, Dale AS, Lunn TJ, Ruiz-Aravena M, Rynda-Apple A, McKee CD, Clark KL. *Borrelia* lineages adjacent to zoonotic clades in black flying foxes (*Pteropus alecto*), Australia, 2018–2020. *Emerg Infect Dis*. 2025 Jul;31(7):1415.

Table S2. Results of generalized additive models for binomial *Borrelia* spp. positivity and tick parasitism in Mexican free-tailed bats ($n$ = 386 and $n$ = 406*, respectively), fit using restricted maximum likelihood through the *mgcv* R package. Predictors are presented with model coefficients or estimated degrees of freedom (EDF) and test statistics. These models explained 4.23% and 17.8% of deviance in *Borrelia* spp. infection and tick parasitism status, respectively.

| | *Borrelia* spp. positivity | | | | | Tick parasitism | | | | |
|---|---|---|---|---|---|---|---|---|---|---|
| Term | OR | $\|z\|$ | EDF | $\chi^2$ | $p$ | OR | $\|z\|$ | EDF | $\chi^2$ | $p$ |
| Intercept | 0.01 | -5.38 | | | <0.001 | 0.01 | -4.31 | | | <0.001 |
| 2023 | 0.24 | -1.15 | | | 0.25 | 11.87 | 2.03 | | | 0.04 |
| Male | 1.62 | 0.39 | | | 0.70 | 0.85 | -0.34 | | | 0.74 |
| s(week) | | | <0.01 | 0 | 0.89 | | | <0.01 | 0 | 0.05 |
| s(week, 2022) | | | <0.01 | 0 | 0.70 | | | 1.62 | 3.16 | 0.14 |
| s(week, 2023) | | | <0.01 | 0 | 0.56 | | | 1.68 | 12.66 | <0.01 |

* One bat of 407 screened individuals lacked data on sex and was thus excluded from the model

Figure S1. Consensus Bayesian phylogeny of the two *flaB Borrelia* spp. sequences from this study (shown in bold) and reference sequences from bats, other mammals, reptiles, and ticks. Nodes are colored by posterior probability (nodes with less than 50% support are not shown).

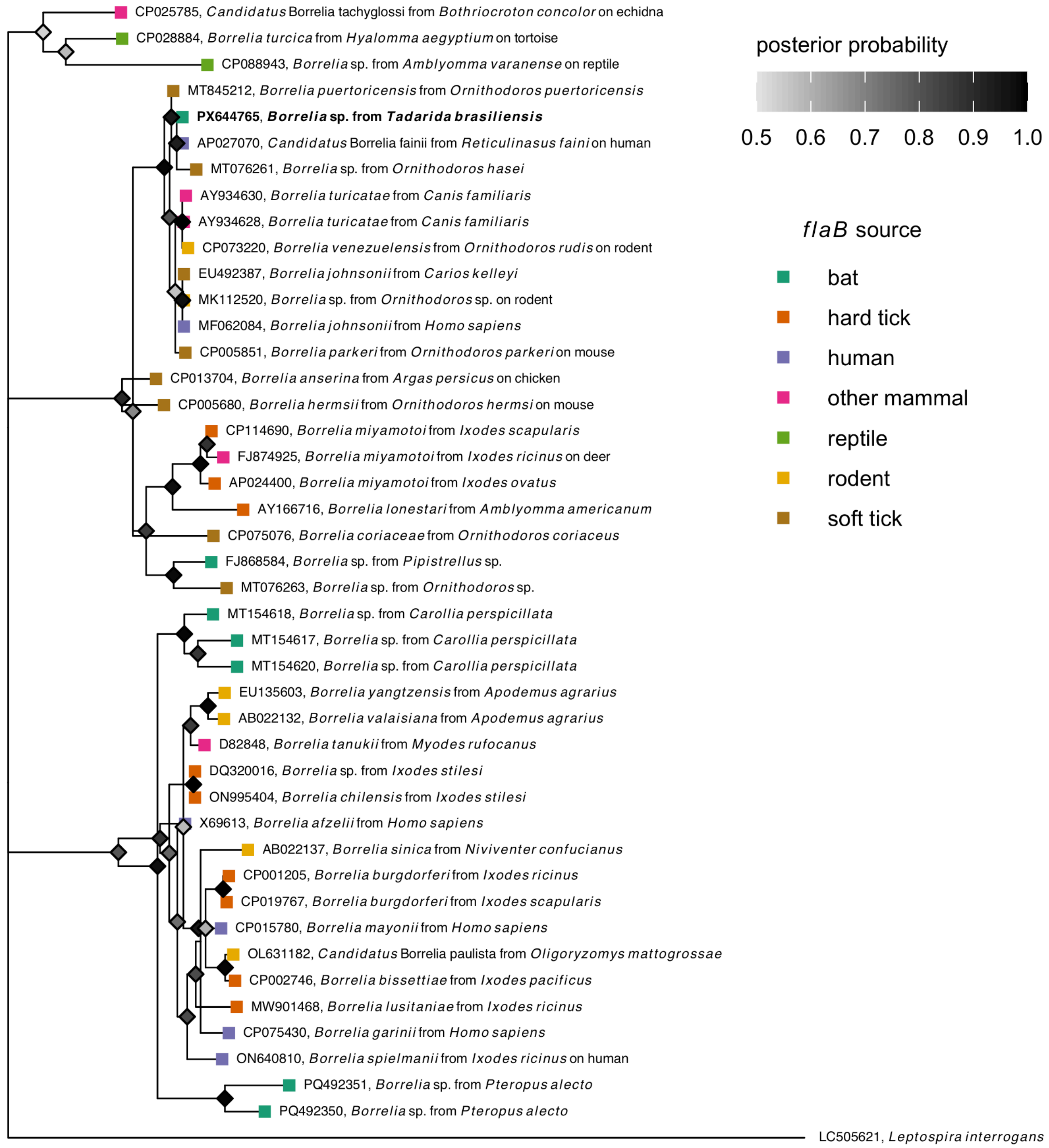

*Additional genomic methods*

**Whole genome amplification and sequencing**. Whole genome amplification was conducted using the TruePrime WGA Kit (1) according to the manufacturer instructions (Expedeon, Heidelberg, Germany).The starting DNA concentration was 0.75 ng DNA in 15 µL. Whole genome amplification yielded 5.64 g DNA. Illumina sequencing was conducted using the services of the Oklahoma State University Next Generation Sequencing Core Facility on an Illumina NextSeq platform using the Watchmaker DNA Library Prep Kit for 600 cycles.

**Bioinformatic analysis**. Two approaches were used simultaneously to probe for the presence of *Borrelia* DNA in the dataset. In the first approach, taxonomic assignment of raw reads was conducted using Kraken 2 (2) as part of the BV-BRC platform (3). Reads identified as affiliated with the genus *Borrelia* were bioinformatically extracted using a custom shell script. The affiliation of these genes to the genus *Borrellia* was confirmed by BLASTn search (4) against the RefSeq database (5).

In the second approach, raw reads were mapped to a suite of 101 genomes belonging to the genus *Borrelia* using Bowtie2 (6). Mapped reads were bioinformatically extracted using SAMtools (7). The affiliation of the extracted reads to the genus *Borrelia* was confirmed using BLASTn (4). The output from both approaches (henceforth OK535_ *Borrelia* reads; File S1) was pooled and dereplicated. We then assembled OK535_ *Borrelia* reads into contigs using three assemblers: MEGAHIT (8), metaSPAdes (9), and IDBA-UD (10). QUAST was used to compare the quality of assemblies (11). Assemblies were then combined and MUMmer was used to obtain a non-redundant set of contigs ($n$ = 401, henceforth combined assembly; File S2) (12). We then queried the combined assembly plus the unmapped fraction of the OK535_ *Borrelia* reads against a *Borrelia*-only protein database (Table S4) using BLASTx to determine the closest relative, percentage amino acid identity, and predicted function of every gene or partial gene fragment recovered (13). Only results with >50 amino acids alignment length were considered.

Table S3. BLAST results identifying the closest relatives, percentage amino acid identity, and gene function of all 622 partial or complete *Borrelia* genes identified. Three different closest relatives were identified, all of which represent different strains of *B. puertoricensis* and exhibit 99.5–99.8% average nucleotide identity (ANI). Strain SUM (GCA_023035875.1) is the type strain of *B. puertoricensis* and was isolated from ticks in central Panama (14); *B. puertoricensis* strain CAU1 (NZ_CP149102) was isolated from ticks collected in Yucatan, Mexico (15); and *B. puertoricensis* strain MN22-0132 (NZ_CP138334.1) genome sequence was submitted by the Mayo Clinic from a human patient. See the attached XLSX file.

Table S4. List of reference *Borrelia* genomes used for read extraction as well as phylogenetic and functional assignment. See the attached XLSX file.

**References**


1. Picher ÁJ, Budeus B, Wafzig O, Krüger C, García-Gómez S, Martínez-Jiménez MI, et al. TruePrime is a novel method for whole-genome amplification from single cells based on TthPrimPol. Nat Commun. 2016 Nov 29;7:13296.

2. Wood DE, Lu J, Langmead B. Improved metagenomic analysis with Kraken 2. Genome Biol. 2019 Nov 28;20(1):257.

3. Olson RD, Assaf R, Brettin T, Conrad N, Cucinell C, Davis JJ, et al. Introducing the Bacterial and Viral Bioinformatics Resource Center (BV-BRC): a resource combining PATRIC, IRD and ViPR. Nucleic Acids Res. 2023 Jan 6;51(D1):D678–89.

4. Altschul SF, Gish W, Miller W, Myers EW, Lipman DJ. Basic local alignment search tool. J Mol Biol. 1990 Oct 5;215(3):403–10.

5. O'Leary NA, Wright MW, Brister JR, Ciufo S, Haddad D, McVeigh R, et al. Reference sequence (RefSeq) database at NCBI: current status, taxonomic expansion, and functional annotation. Nucleic Acids Res. 2016 Jan 4;44(D1):D733–45.

6. Langmead B, Salzberg SL. Fast gapped-read alignment with Bowtie 2. Nature methods. 2012;9(4):357–9.

7. Li H, Handsaker B, Wysoker A, Fennell T, Ruan J, Homer N, et al. The Sequence Alignment/Map format and SAMtools. Bioinformatics. 2009 Aug 15;25(16):2078–9.

8. Li D, Liu CM, Luo R, Sadakane K, Lam TW. MEGAHIT: an ultra-fast single-node solution for large and complex metagenomics assembly via succinct de Bruijn graph. Bioinformatics. 2015 May 15;31(10):1674–6.

9. Nurk S, Meleshko D, Korobeynikov A, Pevzner PA. metaSPAdes: a new versatile metagenomic assembler. Genome Res. 2017 May;27(5):824–34.

10. Peng Y, Leung HCM, Yiu SM, Chin FYL. IDBA-UD: a de novo assembler for single-cell and metagenomic sequencing data with highly uneven depth. Bioinformatics. 2012 Jun 1;28(11):1420–8.

11. Gurevich A, Saveliev V, Vyahhi N, Tesler G. QUAST: quality assessment tool for genome assemblies. Bioinformatics. 2013 Apr 15;29(8):1072–5.

12. Marçais G, Delcher AL, Phillippy AM, Coston R, Salzberg SL, Zimin A. MUMmer4: A fast and versatile genome alignment system. PLoS Comput Biol. 2018 Jan;14(1):e1005944.

13. Gish W, States DJ. Identification of protein coding regions by database similarity search. Nat Genet. 1993 Mar;3(3):266–72.

14. Bermúdez SE, Armstrong BA, Domínguez L, Krishnavajhala A, Kneubehl AR, Gunter SM, et al. Isolation and genetic characterization of a relapsing fever spirochete isolated from

Ornithodoros puertoricensis collected in central Panama. PLoS Negl Trop Dis. 2021 Aug;15(8):e0009642.

15. Vázquez-Guerrero E, Kneubehl AR, Reyes-Solís GC, Machain-Williams C, Krishnavajhala A, Estrada-de Los Santos P, et al. Use of a mouse model for the isolation of Borrelia puertoricensis from soft ticks. PLoS One. 2025 Feb 18;20(2):e0318652.
